# Baseline control of optimal performance in recurrent neural networks

**DOI:** 10.1101/2022.05.11.491436

**Authors:** Shun Ogawa, Francesco Fumarola, Luca Mazzucato

## Abstract

Changes in behavioral state, such as arousal and movements, strongly affect neural activity in sensory areas. Recent evidence suggests that they may be mediated by top-down projections regulating the statistics of baseline input currents to sensory areas, inducing qualitatively different effects across sensory modalities. What are the computational benefits of these baseline modulations? We investigate this question within a brain-inspired framework for reservoir computing, where we vary the quenched baseline inputs to a random neural network. We found that baseline modulations control the dynamical phase of the reservoir network, unlocking a vast repertoire of network phases. We uncover a new zoo of bistable phases exhibiting the simultaneous coexistence of fixed points and chaos, of two fixed points, and of weak and strong chaos. Crucially, we discovered a host of novel phenomena, including noise-driven enhancement of chaos and ergodicity breaking; neural hysteresis, whereby transitions across phase boundary retain the memory of the initial phase. Strikingly, we found that baseline control can achieve optimal performance without any fine tuning of recurrent couplings. In summary, baseline control of network dynamics opens new directions for brain-inspired artificial intelligence and provides a new interpretation for the ubiquitously observed behavioral modulations of cortical activity.

## Introduction

The activity of neurons across cortical areas is strongly modulated by changes in behavioral state such as arousal [1, 2], movements [3, 4, 5, 6], and task-engagement [7]. Intracellular recordings showed that these behavioral modulations are mediated by a change of baseline synaptic currents, likely originating from the thalamus and other subcortical areas [8, 9]. Such baseline modulations exert strong effects on neural activity explaining up to 50% of its variance across cortical areas, a much larger effect compared to the task-related modulations [4, 5, 6].

What are the functional effects of these baseline modulations on brain function? Experimental results across different sensory modalities painted a contradictory picture. Locomotion-induced modulations can improve visual processing [3, 10, 11], but degrade auditory processing [12, 13, 14]. Arousal, measured by pupil size, can improve gustatory and auditory processing at low to intermediate levels [2, 15, 16], but degrades performance auditory processing at high levels [17]. This variety of complex and apparently contradictory effects of behavioral modulations on neural activity and task performance poses a challenge to current theories of brain function and cortical circuit dynamics.

We aim to shed light on the effects of baseline modulations on cortical activity within the framework of reservoir computing, a powerful tool based on recurrent neural networks (RNNs) with random couplings. Random RNNs can recapitulate different dynamical phases observed in cortical circuits, such as silent or chaotic activity [18], fixed points [19], and the balanced regime [20]; and provide a simple explanation for task selectivity features [21] and the heterogeneity of timescales [22] observed in cortical neurons. Random RNNs can achieve optimal performance in memory tasks when poised at a critical point either by fine-tuning their random couplings [23] or their noisy input [24].

Following recent theoretical [2, 11] and experimental studies [4, 25], we modeled the effect of changes in an animal’s behavioral state as changes in the mean and across-neurons variance of the constant baseline input currents to an RNN. We found that baseline modulations steer the network activity to interpolate between a large zoo of dynamical phases. Beyond known phases, such as fixed points and chaos, baseline modulations unlocked new ergodicity-breaking phases, where the network activity can switch between weak and strong chaos, between a fixed point and chaos, or between two fixed points, depending on the initial conditions. All these different phases were continuously connected and achieved without any training or fine tuning of synaptic couplings. Strikingly, we found a new effect where an increase in quenched noise can induce chaos. When interpolating adiabatically between phases via baseline modulations, we uncovered the novel phenomenon of neural hysteresis, whereby the network activity retains a memory of the path followed in phase space.

Our theory further revealed that baseline modulations can flexibly control optimal performance in a sequential memory task at the edge of chaos, without any fine-tuning of synaptic weights. More generally, our theory shows that baseline modulations unlock a much richer dynamical phase portrait for RNNs than previously known. Baseline control represents a very simple and efficient way for a reservoir to flexibly toggle its dynamical regime to achieve different computations. Our results thus suggest a novel computational role for behavioral modulations of neural activity, suggesting that they might allow cortical circuits to flexibly adjust the cognitive task they perform to adapt to different contexts.

## Results

We model our local cortical circuit as a recurrent neuronal network (RNN) of N neurons where the synaptic couplings are randomly drawn from a Gaussian distribution of mean *J*_0_/N and variance *g*^2^/N (Fig. 1A). We choose a positive definite neuronal transfer function *ϕ*(*x*) = 1/[1 + exp(*x* − *θ*_0_)] with threshold *θ*_0_. Every neuron in our model receives a constant external synaptic input b_*i*_ drawn from a Gaussian distribution with mean µ and variance *σ*^2^. This baseline represents the afferent projections to the local cortical circuit originating from other areas. Following experimental [8, 9, 25] and theoretical studies [2, 11], we modeled behavioral modulations as a change in the baseline statistics (mean µ and variance *σ*^2^) of synaptic inputs b_*i*_ to the local circuit, induced by topdown projections carrying information about the behavioral state of the animal, or other contextual modulations [2, 11]. Because the characteristic timescale of behavioral modulations is typically much slower than a circuit’s stimulus responses, we approximate the effects of such modulations as the quenched inputs b_*i*_. Importantly, this baseline modulations are constant, time-independent offsets of the input current to each neurons, and represent *quenched* input noise.

**Fig. 1.**
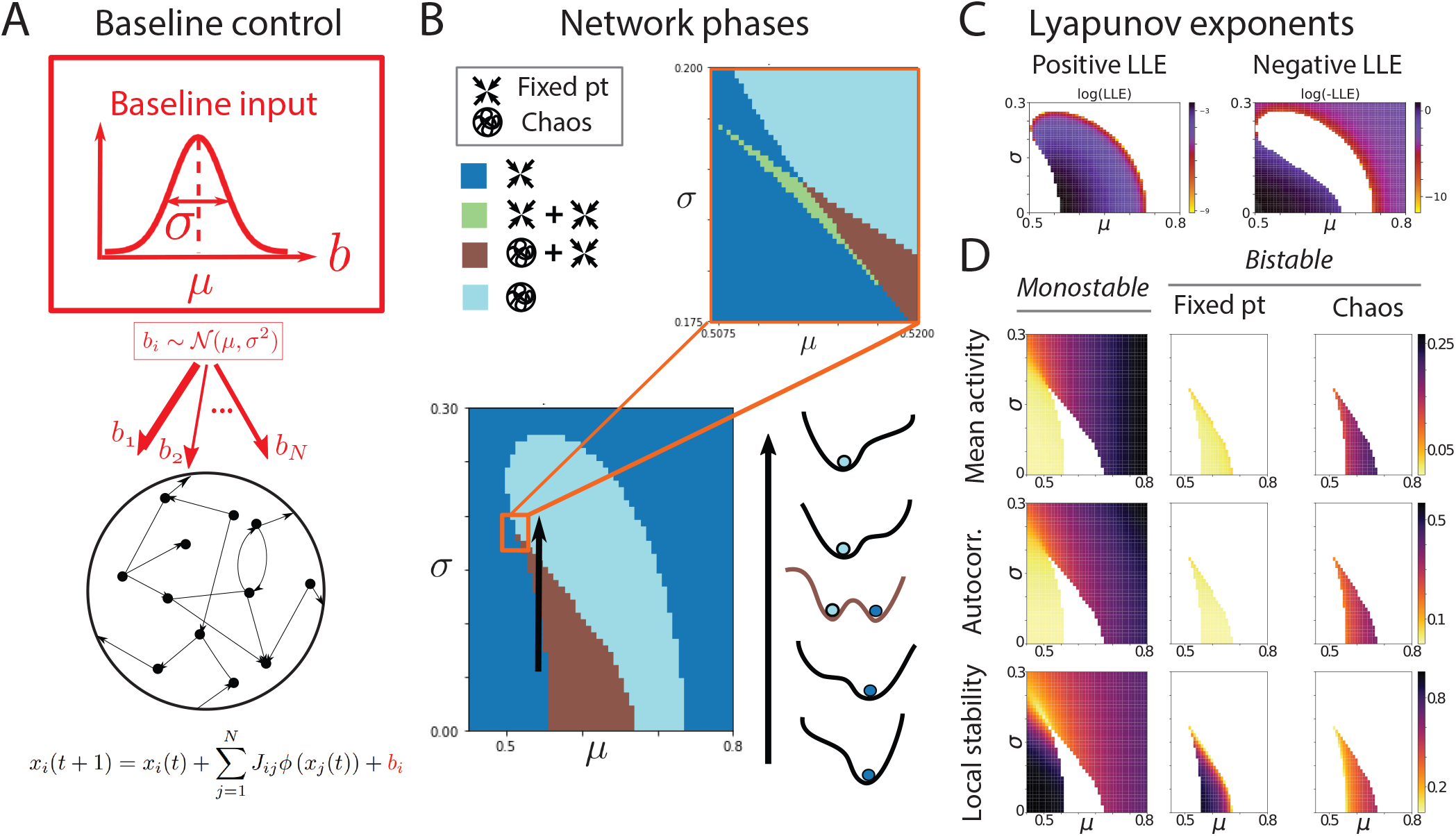
Baseline control of the network dynamical phase. A) Random neural network where the baseline input current b_*i*_ to the i-th neuron is drawn from a normal distribution 𝒩 (*µ, σ*^2^) (red). B) Left: Network phase diagram, obtained by varying the mean µ and variance *σ*^2^ of the baseline input, shows four phases: fixed point (blue); chaos (cyan); bistable phase with coexistence of two fixed points (green); bistable phase with coexistence of fixed point and chaos (brown). Top right: multi-critical point. Bottom right: Schematic of in Landau potential along a phase space trajectory (black arrow in inset) from a stable phase with a single fixed point (blue circle), to a bistable phase with coexistence of fixed point and chaos (blue and cyan circles), to a stable phase with chaos (cyan circle). Positive (left) and negative (right) largest Lyapunov exponents (in the bistable phases both LLE coexist); D) Order parameters in each phase: Mean network activity (top); autocorrelation (middle); local stability (bottom). Representative network activity in the different phases. Insets: Order parameters (Autocovariance *C*_0_, *C*_∞_ and mean activity M). Network parameters: *J*_0_ = 0.5, g = 5, *θ*_0_ = 1.

### Baseline control of the network dynamical phases

We found that by varying the values of baseline mean and variance µ, *σ*^2^, one can access a large library of network phases (Fig. 1b-c). The first two phases are generalizations of the fixed point and the chaotic phase which were previously reported in [26]. Strikingly, we found a zoo of new phases including new ‘bistable’ phases where the network activity can reach two different dynamical branches for the same values of recurrent couplings and baseline input, depending on the initial conditions. In the network of Fig. 1B, the bistable phases are of two different kinds, with coexistence of either a fixed point and chaos (brown) or two fixed points (green). Whereas in the monostable phase the network Landau potential has one global minimum, in the bistable phases it exhibits two local minima, each one defining the basin of attraction of the initial conditions leading to each of the two bistable branches. Depending on the statistics of the random couplings (*J*_0_, *g*), we found networks with up to five different phases, including a new bistable phase featuring the coexistence of strong and weak chaos (see Supplementary Material). Each monostable phase and each branch of a bistable phase can be captured in terms of the network order parameters LLE, *M* and *C* (Fig. 1C), representing, respectively, the largest Lyapunov exponent LLE and the mean *M* and variance *C* of the activity obtained from the self-consistent dynamic mean field equations (see Methods).

The variance of the activity includes a contribution *σ*^2^ from the quenched baseline input and a recurrent contribution. A useful characterization of the network dynamical phase is obtained when considering the population-averaged autocorrelation function *c*(*t*) at lag t; in particular, its zero lag value *c*(0) = *C* the network variance, and its asymptotic value for large lag *c*(∞). The network is at a fixed point if *c*(*t*) does not depend on time (i.e., *c*(∞) = *c*(0) = *C*), while it is in a chaotic phase if c(0) > c(∞), in which case the LLE is positive. Finally, a value of c(∞) > 0 signals a nonzero mean activity driven by the quenched variance in the baseline input.

### Noise-induced enhancement of chaos

Exploring the features of baseline modulations revealed a novel and surprising phenomenon, whereby increasing the variance of the quenched input can enhance chaos. This phenomenon can be understood from a mean field perspective by considering how the baseline and the recurrent synaptic inputs interact with the single cell transfer function to determine the operating point of the network dynamics (Fig. 2). To illustrate this phenomenon, we first revisit the known case of noise-driven suppression of chaos realized in a circuit with quenched inputs and a zero-centered transfer function (Fig. 2A), which can be obtained when the mean baseline is set equal to the threshold µ = *θ*_0_ (see [27] for a case where they both vanish). On general grounds, one expects the network phase to be chaotic whenever a large fraction of the synaptic input distribution is concentrated in the high gain region of the transfer function, in which *ϕ*^′^(*x*)^2^ is large. The distribution of synaptic inputs has mean M, which is centered at the threshold, and some nonzero variance *C*, obtained self-consistently from (3) and (4). For zero baseline variance, the network exhibits chaotic activity (case 1), as a large fraction of the synaptic inputs have access to the high gain region of the transfer function. When turning on a quenched baseline variance *σ*^2^, the synaptic input increases its variance by a value proportional to *σ*^2^. For larger values of the baseline variance *σ*^2^, the fraction of synaptic inputs in the high gain region progressively shrinks and for large enough variance chaos is suppressed (case 2).

**Fig. 2.**
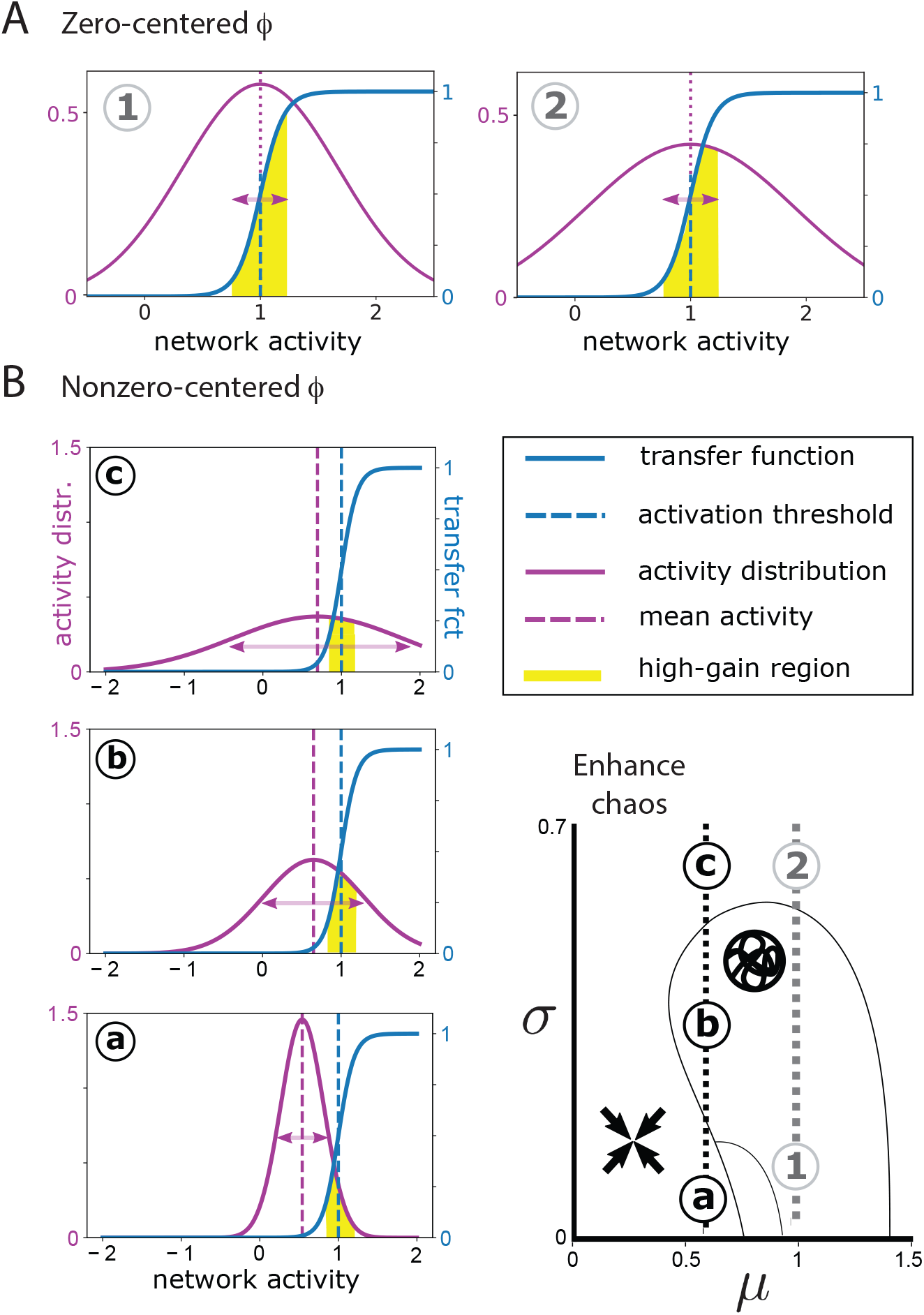
Noise-driven modulations of chaos. A) When the transfer function input is zero-centered (i.e., the baseline mean *µ* equals the threshold *θ*_0_). The chaotic phase (1) occurs when a large fraction of the synaptic input distribution (pink curve) lies within the high gain region (yellow shaded area) of the transfer function (blue curve). Increasing the input quenched variance (2) reduces this fraction, suppressing chaos. B) When the transfer function is nonzero-centered (*µ* < *θ*_0_), for low (a) and high (c) quenched input variance network activity is at a fixed point, because the high gain region receives a small synaptic input fraction; this fraction is maximized at intermediate quenched input variance (b), enhancing chaos. Network parameters: *g* = 5, *θ*_0_ = 1, *J*_0_ = 0. Panel A: *ϕ*(x) = tanh(x − *θ*_0_), *µ* = 1; cases (1, 2) *σ* = (0.1, 0.6). Panel B: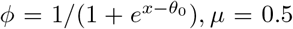; case (a, b, c): *σ* = (0.2, 0.4, 0.9).

In the case where *µ* < *θ*_0_, the transfer function is not zero-centered, and noise-driven enhancement of chaos can occur (Fig. 2B). For low baseline variance *σ*^2^, the network is in the fixed point regime as a small fraction of synaptic inputs has access to the high gain region (case a). Increasing the baseline variance *σ*^2^ leads to a transitions into a chaotic phase, as a progressively larger fraction of synaptic inputs has access to the high gain region. At some large enough variance, though, the fraction of synaptic inputs in the high gain region starts decreasing again and eventually this leads to a new transition to the fixed point phase. This chaos enhancement can be achieved either by passing through an intermediate bistable phase (black arrow in Fig. 1B); or by inducing a direct transition from a fixed point to a chaotic phase at lower values of the mean baseline *µ* (gray arrow). This novel chaos enhancement has a number of striking consequences, such as baseline control of optimal performance and neural hysteresis, which we will examine in the next sections. While previous studies showed that an increase in the *temporal* noise (e.g., white noise inputs) always leads to suppression of chaos [28, 27, 29, 24], we found that quenched noise unlocks a much richer set of phenomena.

### Ergodicity breaking in the bistable phases

The network activity in a bistable phase exhibits dynamical breaking of ergodicity. To illustrate this effect, we consider a network with fixed baseline mean *µ* at different values of *σ* (Fig. 3). At intermediate values of *σ* the network is in the bistable phase featuring a coexistence of a fixed point attractor and chaos, while at low and high values the network is in the monostable fixed point phase and the chaotic phase, respectively. In the bistable phase, the network dynamics converge to either a fixed point attractor or to a chaotic attractor, depending on the initial conditions (Fig. 3A). These two branches are characterized by a negative (fixed point) or a positive (chaos) LLE, respectively, and branch-specific values values for the network order parameters (*C, M*, Fig. 3B). We quantified ergodicity breaking in terms of the average distance ⟨d(T)⟩ between temporal trajectories (starting from different initial conditions, or between different replicas) over an epoch *T* (Fig. 3C). In the ergodic monostable phases ⟨*d*(*T*)⟩ converges to *C*_∞_ at large T → ∞, since the network activity eventually explores all possible configurations (in the chaotic phase, the decay is typically slower than in a phase with a single attractor). If ⟨*d*(*T*)⟩ does not decays to *C*_∞_ but it monotonically increases to reach a non-zero late time values larger than *C*_∞_, the network breaks ergodicity. This means that, depending on the initial conditions, there are different basins at finite distance from each other. We found that the network is non-ergodic in the all the bistable phases, although each one of these phases retains specific values of the order parameters.

**Fig. 3.**
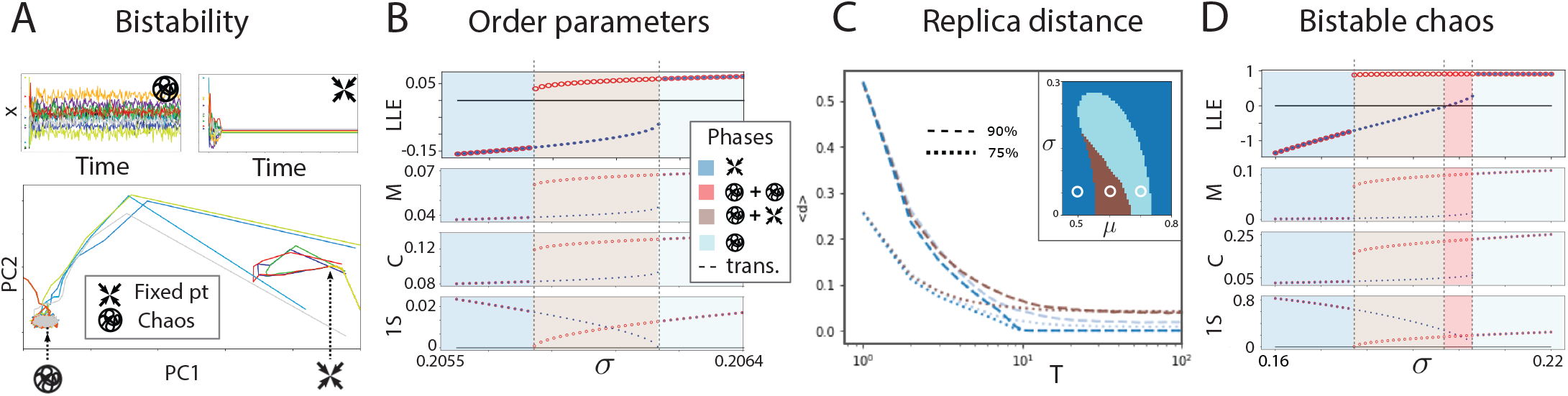
Ergodicity breaking in the bistable phase. A) Representative trajectories of network activity (top) and their top two principal components (bottom) in the bistable phase obtained from initial conditions within the chaotic (left) and fixed point (right) basins of attraction. B) For increasing values of *σ*, a crossover from a monostable fixed point phase (left), to a bistable phase fixed point/chaos (middle) to a monostable chaotic phase (right) is revealed by the order parameters (LLE: Largest Lyapunov exponent; M: mean activity; *C*: mean autocorrelation; 1S: 1-replica stability). In the bistable phase, the fixed point and chaotic branches exhibit different order parameters. C) Average distance between replica trajectories < d > reveals ergodicity breaking: in the monostable fixed point (blue) and chaotic (cyan) phases < d > asymptotes to *C*_∞_, but in the bistable phase (brown) it asymptotes to a value larger than *C*_∞_, representing the average distance between the basins of attraction of the two branches. (D) Example of a crossover from a monostable fixed point phase (blue), to a bistable phase fixed point/chaos (brown) to a bistable weak/strong chaos phase (red), to a monostable chaotic phase (cyan), as revealed by the order parameters (same as panel B). Network parameters: panel A: J_0_ = 0.5, *θ*_0_ = 1, *µ* = 0.54, g = 5, *σ* = 0.1; panel B: J_0_ = 0.5, g = 6, *θ*_0_ = 1, *µ* = 0.5; panel *C*: same as Fig. (1B) and *µ* = 0.5, 0.6, 0.7; panel D: J_0_ = 0.5, *θ* = 1, *µ* = 0.5, g = 18.

The library of bistable phases induced by changes in the baseline statistics include all the phases in Fig. 3 and, remarkably, a novel phase exhibiting the coexistence of two chaotic phases 3D. This double chaos phase features a weak chaotic branch with small positive LLE and slow dynamics, and a strong chaotic branch with large positive LLE and fast dynamics. We found that this double chaos phase occurs for large g and it exhibits important computational properties that we investigate below.

### Neural hysteresis retains memory of network phase trajectories

What are the effects of adiabatic changes in baseline statistics on the network dynamics? We sought to elucidate the effects of slow baseline changes, by driving the network with time-varying values of *σ*(*t*) for fixed *µ*, describing a closed loop (Fig. 4). We found that the network order parameters *C, M* changed discontinuously across phase boundaries, signaling a phase transition. When the baseline trajectory crosses the phase boundary from a stable phase (with a single LLE) to a bistable phase (with two branches, each characterized by its own LLE), the network activity in the bistable phase lies on either of the two branches, characterized by two separates basins of attractions (Fig. 3). The rules governing which of the two branches will be reached are determined by a novel hysteresis effect. We found that the network activity in the bistable phase retained a memory of the dynamical branch that it occupied before crossing the phase boundary. In the particular example of Fig. 4, when crossing the boundary from the monostable fixed point to the bistable phase, the activity will persist on the fixed point branch of the bistable phase, whose negative LLE is continuously connected with the fixed point phase. For larger values of *σ*(*t*), the network will eventually enter the monostable chaos phase, where the LLE discontinuously jumps to a very large value. Vice versa, when inverting the time-varying trajectory in phase space by slowly decreasing the *σ*(*t*) from the monostable chaotic phase into the bistable phase, the network will persist on the chaotic branch of the latter, whose positive LLE is continuously connected to the monostable chaotic phase. Eventually, for lower *σ*(*t*) the network falls back into the fixed point phase where the LLE discontinuously jumps from large positive to negative values. Thus, when crossing phase boundaries adiabatically the network will choose the branch of the bistable phase whose LLE is continuously connected to the previous phase.

**Fig. 4.**
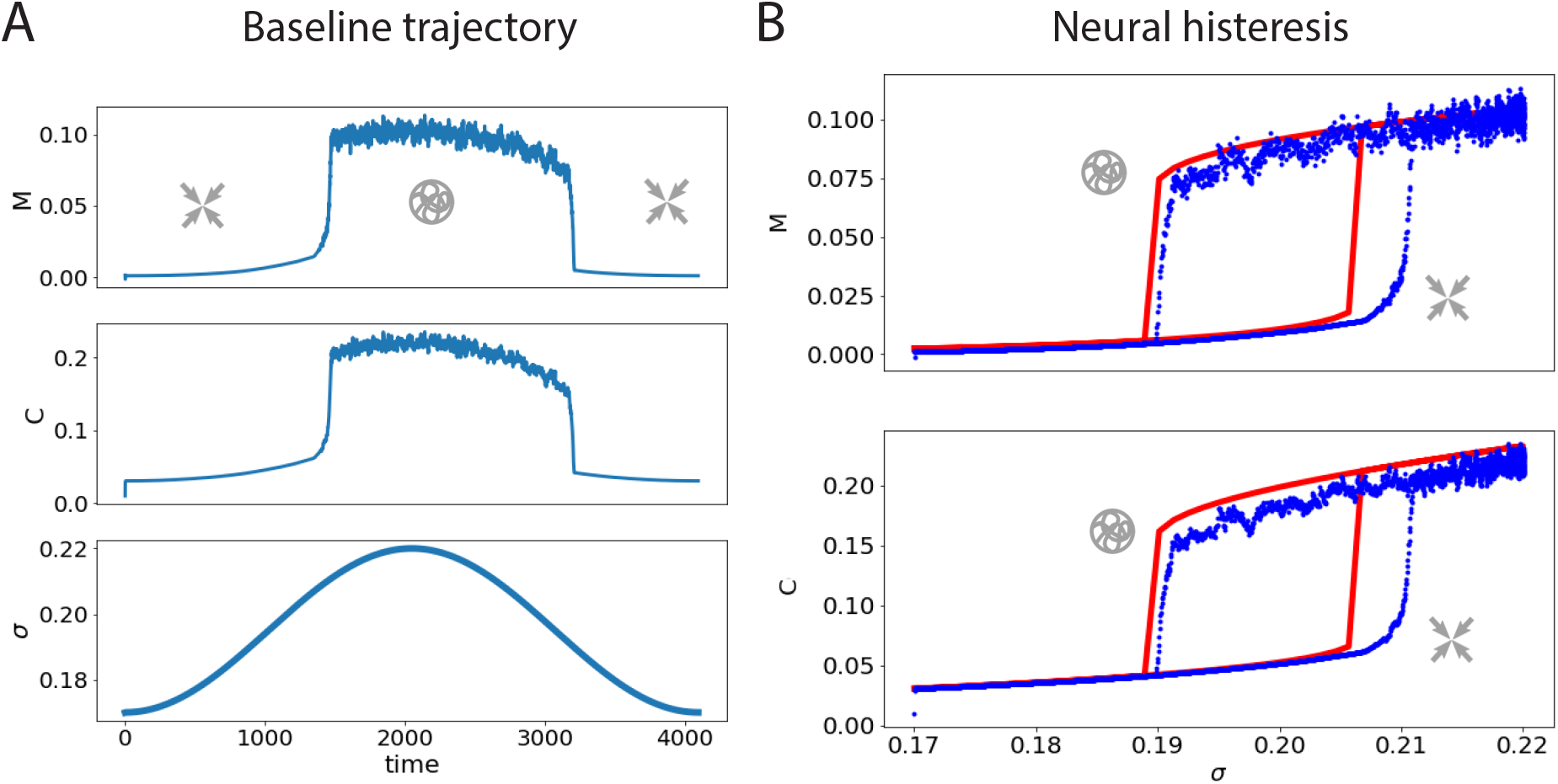
Neural hysteresis. A) Slow changes in baseline variance *σ*(*t*) leads to discontinuous transitions in the network order parameters M, *C* (left: temporal profile of *M, C, σ*). B) Crossing over phase boundaries by a time-varying *σ*(*t*) retains memory of the previous phase (blue: network simulations; red: exact DMFT calculation). Network parameters: J_0_ = 0.5, g = 12, *θ*_0_ = 1, *µ* = 0.5, *σ*(*t*) = *σ*_0_ + *σ*_1_ sin(*πt/T*) with *T* = 2048, *σ*_0_ = 0.17, *σ*_1_ = 0.025.

Neural hysteresis occurs not just in the fixed point/chaos bistable phase, but also in the double fixed point and double chaos bistable phases. When crossing boundaries between two adjacent bistable phases, more complex hysteresis profiles can occur. For example, when crossing into the double chaos phase (with fast/slow chaotic branches, Fig. 3D), from the fixed point branch of the fixed point/chaotic phase, the network dynamics will lie on the slow chaotic branch, whose positive but small LLE is continuously connected to the fixed point branch of the previous bistable phase. However, when crossing into the double chaos phase from the chaotic branch of the fixed point/chaotic bistable phase, the network dynamics will persist on the fast chaotic branch, whose large positive LLE is continuously connected to the chaotic branch of the fixed point/chaotic bistable phase. We then examinedthe relevance of neural hysteresis for controlling the network performance in a memory task.

### Baseline control of optimal memory capacity

A classic result in the theory of random neural networks is that, by fine tuning the recurrent couplings at the ‘edge of chaos’, one can achieve optimal performance in a memory task, where the network activity maintains for a very long time a memory of stimuli presented sequentially [23]. This was achieved by fine tuning the network recurrent couplings to values close to the transition between fixed point and chaos, which is a metabolically costly and slow procedure typically requiring synaptic plasticity. Is it possible to achieve optimal memory capacity without changing the recurrent couplings? We found that baseline control can achieve optimal memory capacity by simply adjusting the mean and variance of the baseline input distribution, without requiring any change in the recurrent couplings, (Fig. 5).

**Fig. 5.**
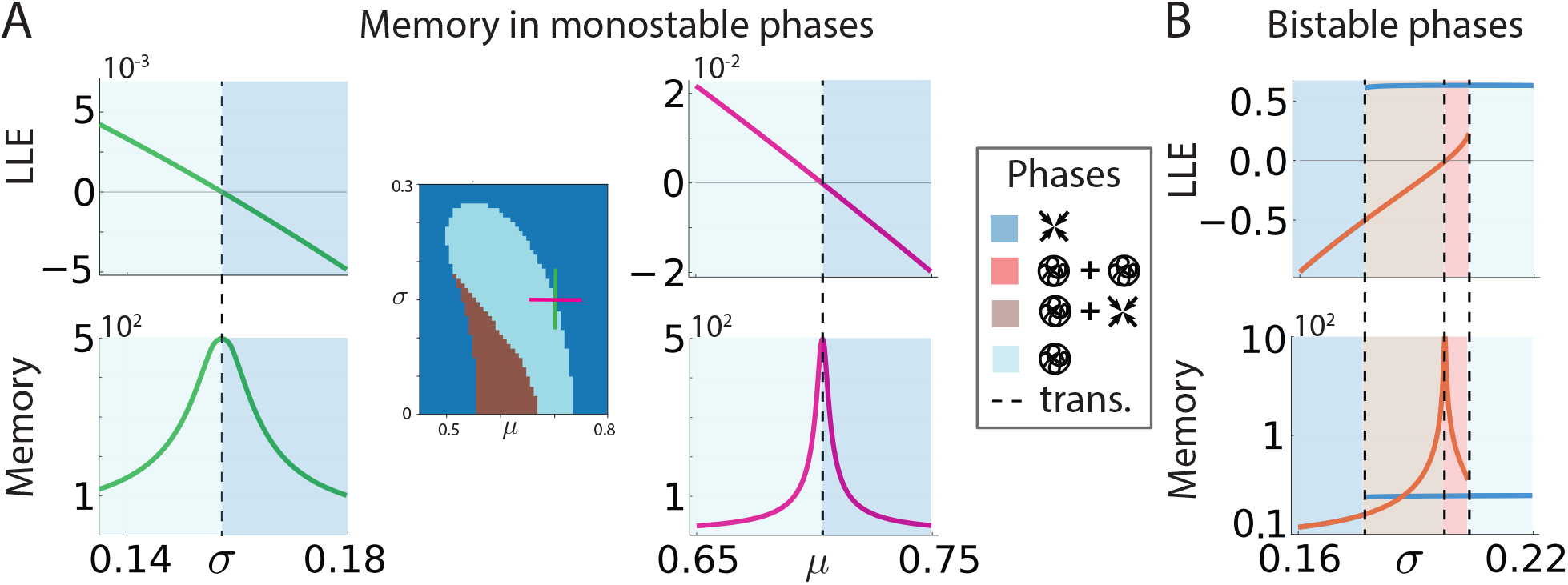
Baseline control of optimal memory capacity. A) Two representative trajectories in baseline (*µ, σ*) space (left: green and orange lines) allow to reach a phase transition where the LLE crosses zero (top panels) and memory capacity is optimized (bottom panels). B) In a transition between bistable phases, memory capacity is optimized by a baseline trajectory whose branch exhibits an LLE that crosses zero at the phase boundary (orange curve); the branch with positive LLE (blue curve) does not maximize memory capacity. Network parameters: Panel A, same as Fig. 1A; panel B, same as Fig. 3D.

We first derived an analytical formula for the memory capacity in the vicinity of a second-order phase transition boundary

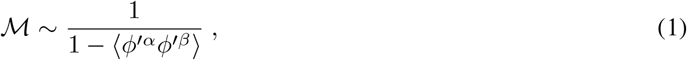

where α, β are replica indices. Optimal memory capacity is achieved close to a phase boundary, and its features are qualitatively different depending on whether the phases separated by the boundary are monostable or bistable. At a boundary between two monostable phases, where the activity transitions between a fixed point and chaotic phase, optimal memory capacity is achieved at the edge of chaos. For fixed values of the recurrent couplings (Fig. 5A), one can easily achieve optimal memory capacity by adiabatically changing either the mean or the variance of the baseline. This external modulation thus sets the network at the edge of chaos, in the region where memory capacity is maximized, via baseline control, without any change in the recurrent couplings. Around a phase boundary involving a bistable phase, the optimal performance region can be reached by making use of the neural hysteresis phenomenon. We illustrate this intriguing scenario in the case of the transition from a bistable fixed point/chaos branch to a bistable double chaos branch (Fig. 5B). Optimal performance is achieved only on the branch of the bistable phase transition which undergoes a second-order phase transition (i.e., the branch whose LLE crosses zero). In this specific case, then, we can reach optimal performance on the lower branch of the LLE curve, describing the transition between the weak chaotic branch of the double chaos phase to the fixed point branch of the fixed point/chaos phase. Because of the neural hysteresis, achieving the optimal performance region requires first initializing the network on the lower LLE branch (on either side of the transition), and then adiabatically controlling the baseline to reach the desired point. The phase boundaries where only first-order phase transitions occur (i.e., no branch exhibits an LLE that crosses zero) do not lead to optimal memory capacity. For example, in Fig. 3B, neither the upper nor lower branch of the transition between a monostable fixed point phase to a bistable fixed point/chaos phase lead to large memory capacity, since no LLE on either branch of the intermediate bistable phase crosses zero. Nevertheless, it is always possible to reach a different second-order phase boundary from any point in (*µ, σ*) space by following an appropriate adiabatic trajectory in the baseline, where optimal memory capacity can be achieved (see Fig. 5A). Therefore, one can achieve baseline control of optimal performance via neural hysteresis.

## Discussion

We presented a new brain-inspired framework for reservoir computing where we controlled the dynamical phase of a recurrent neural network by modulating the mean and quenched variance of its baseline inputs. Baseline modulations revealed a host of new phenomena. First, we found that they can set the operating point of the network activity by controlling whether synaptic inputs overlap with the high gain region of the transfer function. A manifestation of this effect is a novel noise-induced enhancement of chaos. Second, baseline modulations unlocked access to a large repertoire of network phases. On top of the known fixed point and chaotic ones, we uncovered a new zoo of bistable phases, where the network activity breaks ergodicity and exhibits the simultaneous coexistence of a fixed point and chaos, of two different fixed points, and weak and strong chaos. By driving the network with adiabatic changes in the baseline statistics one can toggle between the different phases, charting a trajectory in phase space. These trajectories exhibited the new phenomenon of neural hysteresis, whereby adiabatic transitions across a phase boundary retain the memory of the adiabatic trajectory. Finally, we showed that baseline control can achieve optimal performance in a memory task at a second-order phase boundary without any fine tuning of the network recurrent couplings.

### Noise-induced enhancement of chaos

Previous theoretical work found a noise-induced suppression of chaos in random neural networks driven by time-varying inputs both in discrete time [28] and continuous time [27, 29, 24, 22, 30, 24]. In previous cases, featuring a mean synaptic input centered in the middle of the high-gain region of the transfer function, suppression of chaos occurs because an increase in the variance drives the network away from the chaotic regime. In contrast, we found that, when the baseline statistics sets the mean synaptic input away from the center of the high gain region, one can induce a transition from fixed point to chaos at intermediate values of the variance (Fig. 2). Larger values of the variance eventually suppress chaos, such that a non-monotonic dependence of the Lyapunov exponent on the baseline variance or mean can be realized. To our knowledge this is the first example of noise-induced chaos in a recurrent neural network (although for the logistic map see [31]). We believe that noise-induced modulation of chaos in discrete time networks is similar for both quenched and dynamical noise [24], since the LLE and the edge of chaos are the same for both cases. We speculate that introducing a leak term and generalizing our results to a continuous time system may induce a dynamical suppression of chaos on general grounds, based on the memory effect. Another interesting direction is to drive the network with dynamical noise at different values of the baseline input and investigate its effect on the different monostable and bistable phases we uncovered via baseline modulation.

### Optimal sequential memory

Previous studies showed that optimal performance in random networks can be achieved by either tuning the recurrent couplings at the edge of chaos [23], by driving the network with noisy input tuned to a particular amplitude [24]. The former method represents a metabolically costly and slow process requiring synaptic plasticity. The latter method may lack biological plausibility, since in a spiking circuit the dynamical input noise statistics are self-consistently determined by the spiking dynamics and are not a tunable parameter. We found that optimal sequential memory performance can be achieved by simple regulation of the mean and variance of the baseline current. Achieving optimal performance by changing the across-neuron variance of baseline currents is a simple and biologically plausible mechanism.

### Information processing capabilities and bistability

Bistable phases in recurrent networks with random couplings were previously reported in [32]. We generalized this to a new set of bistable phases featuring the coexistence of two fixed points and, remarkably, two chaotic attractors with slow and fast chaos, respectively. To our knowledge, this is the first report of a doubly chaotic phase in recurrent neural networks. Are there any information processing benefits of the double chaos phase? Neural activity unfolding within the weakly chaotic branch of this bistable phase has large sequential memory capacity, as the Fisher information diverges at the edge of chaos. On the other hand, the strongly chaotic branch erases memory fast. In this doubly chaotic phase, the network’s information processing ability can be changed drastically by switching between the two branches, for example via an external pulse. It would be tantalizing to explore the computational capabilities of these new bistable phases unlocked by baseline modulation. Here, we only considered homogeneous inputs where the baseline statistics is the same for all network neurons. Although, one may consider a more general set up with heterogeneous inputs, where different neural populations receive baseline modulations with different statistics. The simplest such possibility would be the ability to perform different tasks by gating in and out specific subpopulations, driving them with negative input. This is a promising new direction for multitasking and we leave it for future work.

### Evidence for baseline modulations in brain circuits

In biologically plausible models of cortical circuits based on spiking networks, it was previously shown that increasing the baseline quenched variance leads to improved performance. This mechanism was shown to explain the improvement of sensory processing observed in visual during locomotion [11] and in gustatory cortex with general expectation [2]. In these studies, the effect of locomotion or expectation was modeled as a change in the constant baseline input to each neuron realizing an increase in the input quenched variance. This model was consistent with the physiological observation of the heterogeneous neuronal responses to changes in behavioral state, comprising a mix of enhanced and suppressed firing rate responses (during locomotion [3, 11, 25], movements [4, 6, 5], or expectation [33, 34]. Intracellular recordings showed that these modulations are mediated by a change of baseline synaptic currents, likely originating from subcortical areas [8, 9]. Because the effects of these changes in behavioral state on neural activity unfolded over a slower timescale (a few seconds) compared to the typical information processing speed in neural circuits (sub-second), we modeled them as constant baseline changes, captured by changes in the mean and variance of the distribution of input currents. Our results provide a new interpretation of these phenomena, leading to the hypothesis that they could enable cortical circuits to adapt their operating regimes to changing demands.

### Neural hysteresis

A new prediction of our model is that baseline modulations may induce neural hysteresis when crossing a bistable phase boundary. Hysteresis is a universal phenomenon observed in many domains of physics. Our results suggest a potential way to examine the existence of hysteresis in brain circuits, within the assumption that increasing baseline variance represents increasing values of a continuous behavioral modulation such as arousal (e.g., measure by pupil size [17]). A potential signature of hysteresis could be detected if the autocorrelation time of neural activity at a specific arousal level exhibited a strong dependence on whether arousal levels decreased from very high levels or increased from very low levels. We leave this interesting direction for future work.

## Methods

### Random neural network model

Our discrete time neural network model with top down control, illustrated in Fig. 1, is governed by the dynamical equation

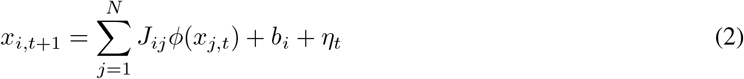

Here b_*i*_ is quenched Gaussian noise with mean *µ* and variance *σ*^2^, η_*t*_ is a possible time-dependent external stimulus (relevant for the sequential memory task below). The mean of the synaptic strength, *J*_0_/N, is not zero and its variance is *g*^2^/N; the scaling 1/N guarantees the existence of the large N limit. We will assume *µ* > 0 in accordance with the fact that top-down modulation is directly conveyed by long-range pyramidal connections. The activation function 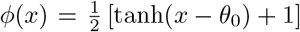 is positive definite and biologically plausible as it incorporates both a soft rectification and thresholding. Indeed the activation function *ϕ* satisfies *ϕ*(x) ≈ 0 when *x* ≪ *θ*_0_ and *ϕ*(*x*) ≈ 1 when *x* ≫ *θ*_0_.

### Order parameters

The order parameters of the model are the population mean and variance at equilibrium of the single neuron activity ⟨*x*_*i,t*_⟩. A rigorous derivation of self-consistent equations for these two quantities requires Dynamical Mean Field Theory (see Supplementary Material), a heuristic argument for them can be sketched as follows. Averaging Eq. 2 in the absence of external input yields

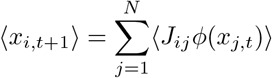

Neglecting correlation between the random variables *J*_*ij*_ and *x*_*j,t*_ on the right hand side, and using the statistical invariance under permutation of neuron labels to drop cell indices, we obtain ⟨*x*_*t*+1_⟩ = *J*⟨*ϕ*(*x*_*t*_)⟩. Focusing now on the stationary regime, where the distribution of *x*_*t*+1_ and *x*_*t*_ are identical, and assumings them to be gaussian with mean *M* and variance *C*, leads to

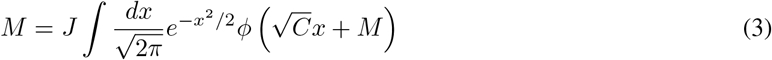

Taking the second moment of Eq. 2, without neglecting the variance of the quenched disorder, term and deploying once again the same assumptions yields

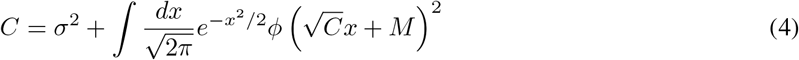

In supplement, the Dynamical Mean-Field Theory approach is rigorously developed to derive two dynamical equations for the mean-field momenta. The stationary limit of those equation is found to correspond to Eqs. 4 and 3, thus confirming the heuristic result.

### Distance between replicas

Let us define the mean activity in the replica a (corresponding to some initial conditions 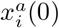) as

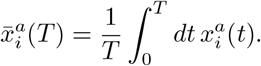

We then define the distance between replicas as [35]

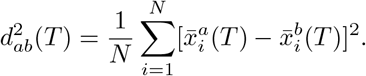

as used in the visualization of Fig. 3C.

### Memory capacity

Following [36, 37], we define the memory capacity of a dynamical system for an observer in possession of an unbiased estimator for the mean, who can therefore remove the mean values from all the time series he records. Moreover, we would like the resulting memory capacity to be zero when the linear readout is dominated by a constant baseline value, because nothing can be learned from a readout independent on the input. Adopting therefore the mean-removed formula, we find for the memory capacity *M* in the neighborhood of the second-order phase transition boundary

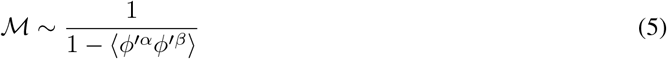

The rigorous derivation of this formula is detailed in Supplement.

### Largest Lyapunov exponent

The Lyapunov exponent of a dynamical system is a quantity that characterizes the rate of separation of infinitesimally close trajectories. Quantitatively, two trajectories in phase space with an initial separation vector diverge (provided that the divergence can be treated within the linearized approximation) at an exponential rate given, and the Lyapunov exponent governs this exponential growth. The LLE for a discrete-time dynamical system can be written as

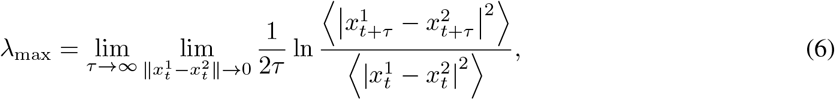

which indicates how the two orbits get to be far from each other.

Going back to the N body picture, we have

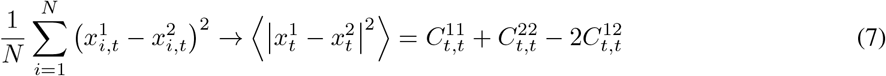

for *N* → ∞. Around the stationary solution, we consider 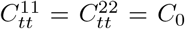 and 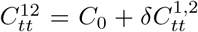. Then we have the LLE as follows;

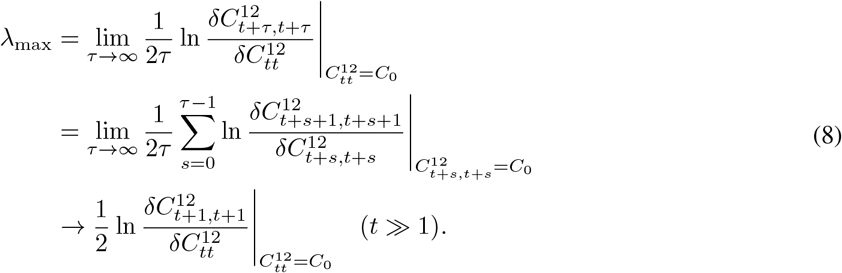

and the LLE is estimated as [38]

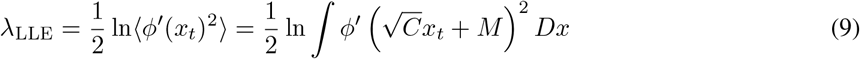

Here *C* and *M* are the stationary solutions *M*_*t*_ and 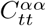 the dynamical mean-field equation in the Supplemental, which are easy to find numerically by iterating substitution. To detect a state of the system (2), what we have to do is just solving Eq. 3 and 4 and check the sign of the LLE (9) for each state. Conceptually, the consequences of Eq. 9 are described in the cartoon of Figure 2. The top-down control can use two levers – mean and variance of its modulation, and depending on the mean, the variance can have the opposite effects of tuning the controlled network into chaos or out of it.

### Contributions and acknowledgments

LM supervised the project; SO worked out the analytics with FF’s support; numerical simulations were carried out by FF and SO; all authors wrote the manuscript. We would like to thank Enrico Rinaldi for advice on the numerics and Taro Toyoizumi for discussions. SO and FF were partially supported by RIKEN Center for Brain Science; LM was supported by National Institute of Neurological Disorders and Stroke grant R01-NS118461 and by National Institute on Drug Abuse grant R01-DA055439 (CRCNS).

## Supplementary Notes

### S1 Dynamical Mean Field Theory

We study the model

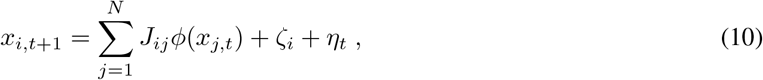

where, as stated in the main text, *x*_*i,t*_ is the individual neuronal activity at time *t, ϕ*(*x*) is the transfer function, ζ_*i*_ is quenched Gaussian noise with mean *µ* and variance *σ*^2^, η_*t*_ is a possible time-dependent external stimulus. The synaptic weights *J*_*ij*_ are randomly drawn from a Gaussian distribution of mean *J*_0_/*N* and variance *g*^2^/*N*.

For this model, the measure of the path integral is

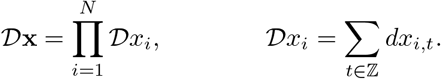

We apply dynamical mean field theory (DMFT) as described in Ref. [39]. The aim of DMFT is to obtain the single body density functional P_1_(*x*) or equivalently its moment generating functional, averaged over the randomness of the synaptic connections and the external noise in the infinite population limit N → ∞. That is,

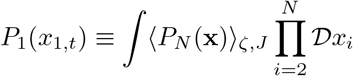

where P_*N*_ (**x**) 𝒟**x** is the N-body density functional. Calling *x*_*i,t*_[*ζ*] the solution to the equations of motion (2) for a given modulation *ζ*, we have

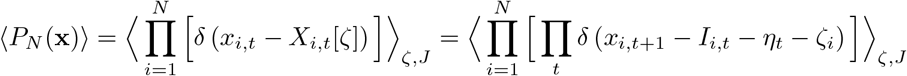

where 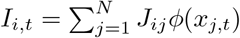 and we changed variables in the path integral noticing that the relevant Jacobian is equal to unity.

Let us now compute the generating functional *Z*_*N*_ [l] over multiple trials or replicas α, written as a function of a control field l:

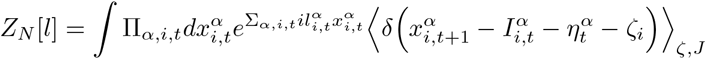

We express the delta function as a Fourier transform, perform the Gaussian integral over the modulation vectors *ζ*, proceed with standard path integral manipulations, and define

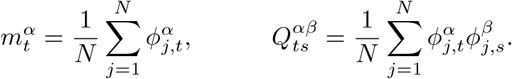

Taking the saddle point in the limit *N* → ∞, we thus obtain a single body generating functional 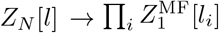, where MF stands for “mean field”:

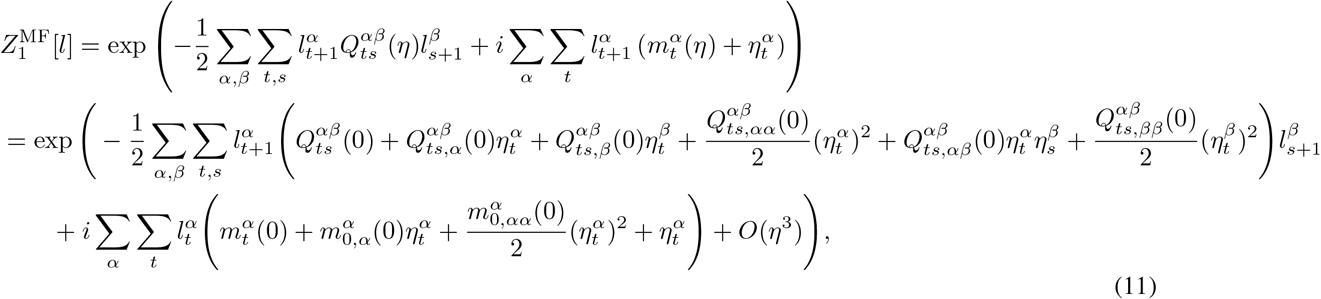

where the subscripts (, *α*) and (, *αβ*) are respectively 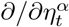 and 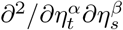; for instance, we have

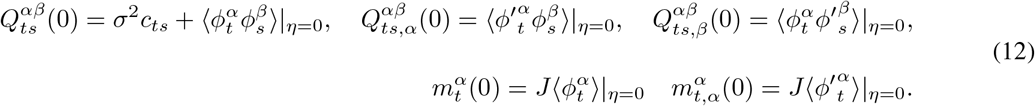

In terms of the generating functional, we finally obtain self-consistent equations for the parameters

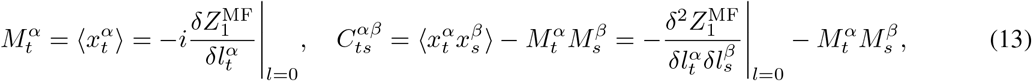

which are explicitly written as follows,

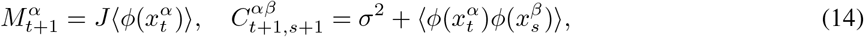

where the indices *α, β* differentiate the individual replicas.

The terms ⟨*ϕ*(*x*_*t*_)⟩ and ⟨*ϕ*(*x*_*t*_)*ϕ*(*x*_*s*_)⟩ are explicitly written as

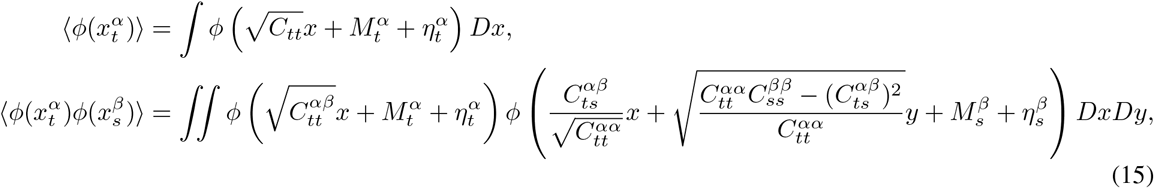

where 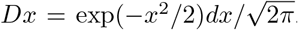. This is because {*x*_*t*_}_*t*∈ 𝕫_ is shown to be a Gaussian random variable whose covariance and mean value is determined self-consistently by use of the generating functional method and by taking mean-field limit *N* → ∞.

From Eqs. 14, we derive

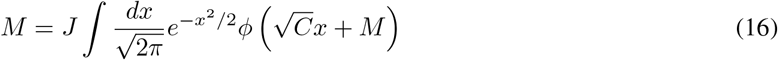

for the fixed point 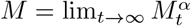 and

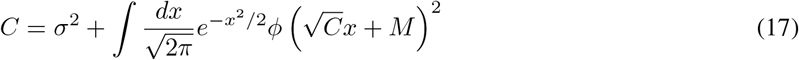

for 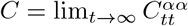. The inter-replica correlation 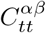 can also be written from the above. Finally, the response to the external force *η* can be computed systematically as

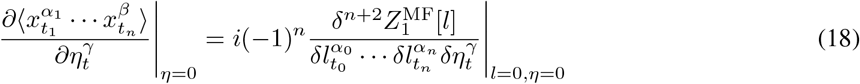

in the infinite population limit.

### S2 Heuristic derivation of conditions for stability

For arbitrary functions *ϕ* and ψ, we define

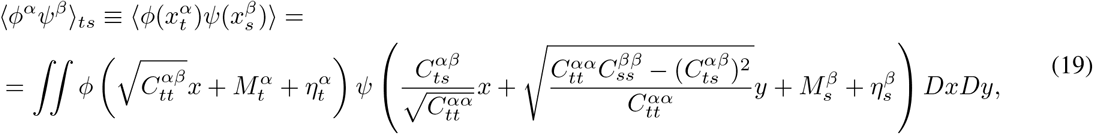

with 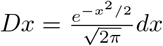

It is easy to see (through integration by parts) that the variation of this quantity under perturbations of *M* ^*α*^ and *C*^*α*^*α* is (omitting time labels for brevity)

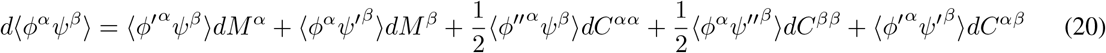

The single-replica stability is understood as follows. Using identity 20 for the quantity ⟨*ϕ*ψ⟩_0_ = ⟨*ϕ*^*α*^ψ^*α*^⟩_*t*_, it is seen that the linearized version of the single-replica equation around the steady state *C*_*tt*_ = *C*^*αα*^, M_*t*_ = *M* ^*α*^, becomes

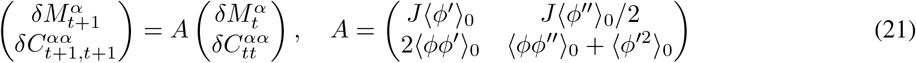

It follows that the steady state is stable if the eigenvalues of A are in the unit circle.

From the above it is also possible to check the stability within one replica, yielding equations for the phase boundaries. The condition of the critical state, *C*_*tt*_ = *C* and *M*_*t*_ = *M*, is indeed

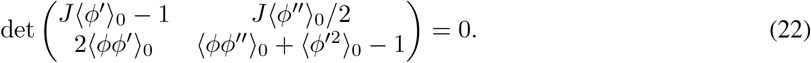

This criticality found within single-replica is on the edge of the coexistence region, not at the edge of chaos in general. In the systems dealt with in Refs. [40, 18, 23], this criticality appears on the edge of chaos due to the symmetry and absence of random noise.

We next consider the stability against the inter-replica perturbation. Invoking once again identity 20, the linearized equation here is found to be

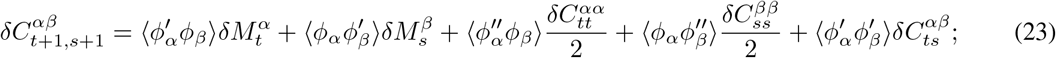

assuming that the system is stable against the intra-replica perturbations 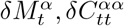 these perturbations converge to 0 so that the linearized equation asymptotically is

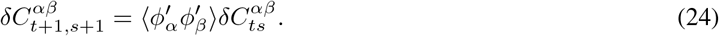

Summarizing the above discussion, the steady state is stable if and only if the eigenvalues of the matrix A are in unit-circle and the inequality 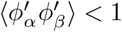 holds.

The single-replica stability is visualized for a range of model parameters in the lowest panels of Fig. 1.

### S3 Field theoretical stability analysis

The stability analysis can also be performed by checking the definiteness of the Hessian matrix around the saddle point [18, 41], e.g. along the lines of Ref. [18]. We will use the abbreviation

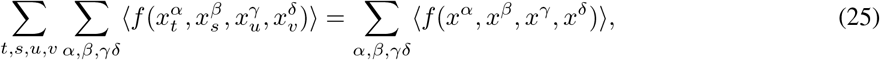

that is, we do not write time parameters (t, s, …) explicitly unless it is necessary, and we only write explicitly the replica parameters as represented by Greek characters. In addition, we will abbreviate 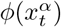 by *ϕ*^*α*^.

With this notation, the generating functional is

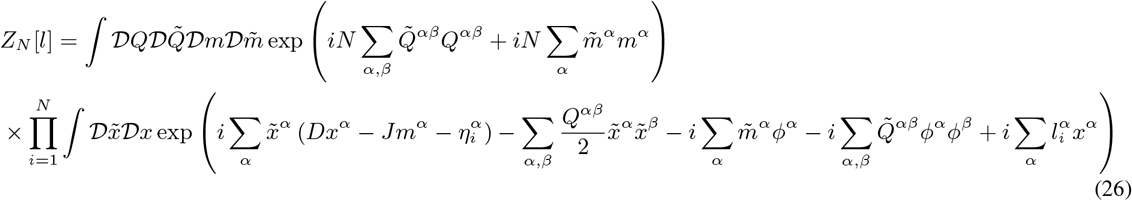

where 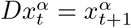.

Let us expand it around the saddle point 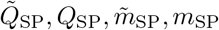 with respect to the fluctuations 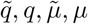 and take the 2nd-order variation.

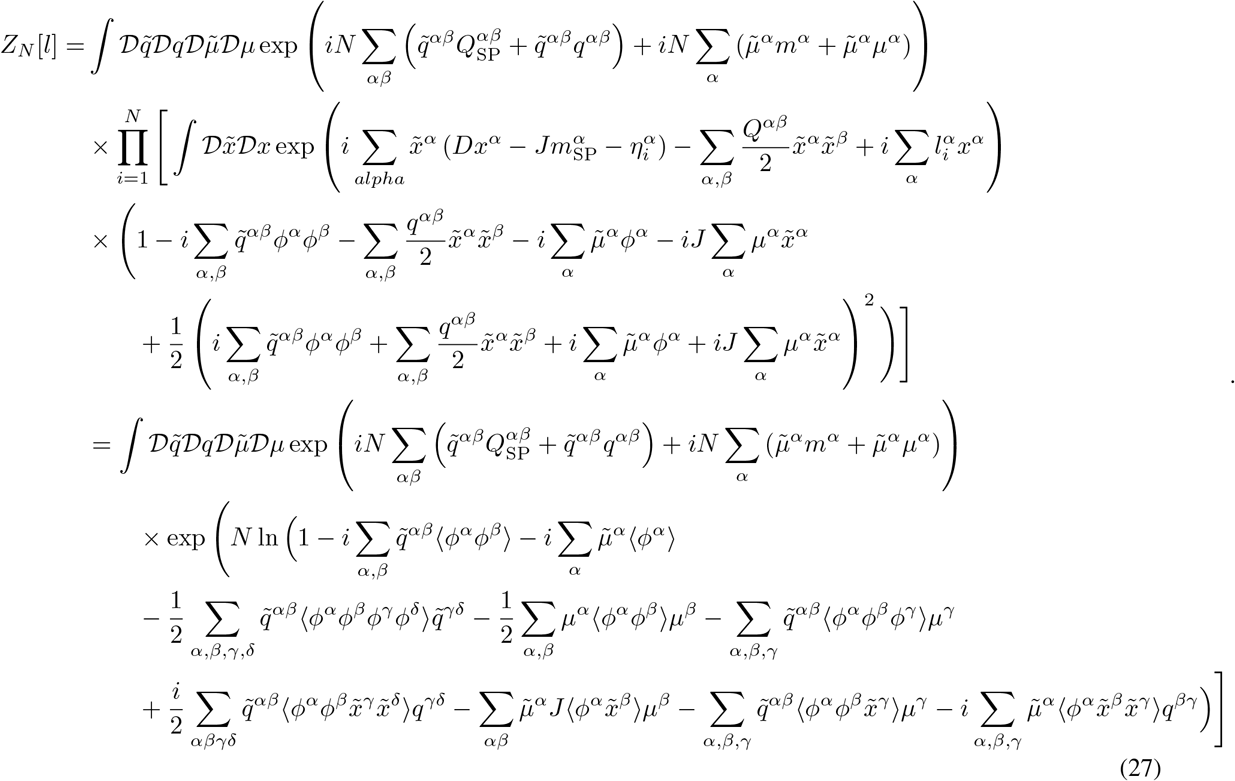

It should be noted that i-dependence coming from η_*i*_ and l_*i*_ is included in the average ⟨•⟩, which may otherwise seems to vanish in the last line of this equation. Using the expansion formula ln 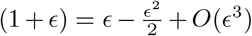, omitting the 3rd order of fluctuations, and using the saddle point condition, we have the 2nd order variation around the saddle point,

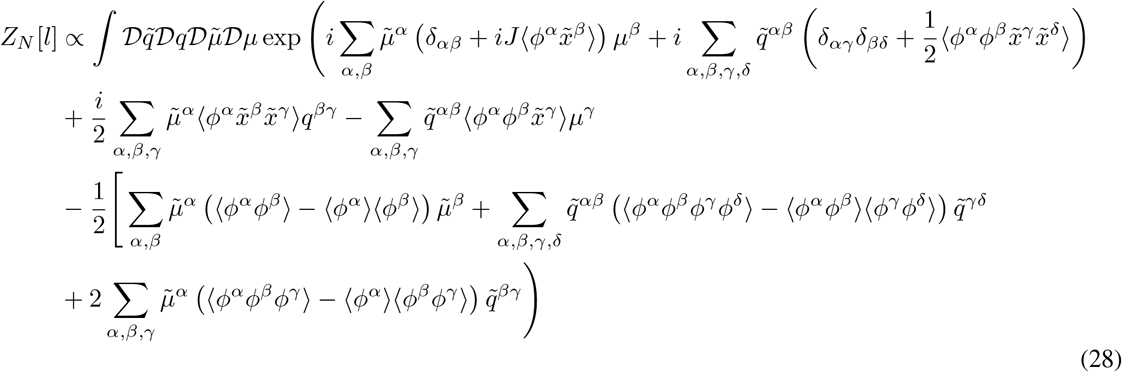

Let us now define the vectors, 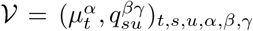 and 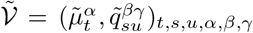. Moreover, let the matrix ℳbe

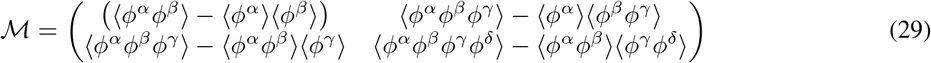

and the matrix 𝒜 be

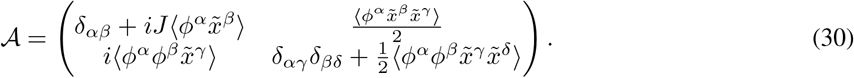

By using them Eq. (28) can be written as

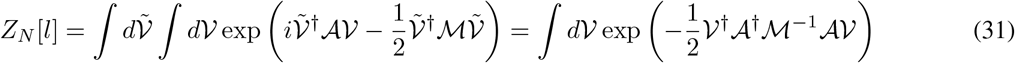

The matrix ℳ is obviously positive definite because it is a covariance matrix. The second variation around the saddle point is thus positive definite if and only if the operator A has no vanishing eigenvalue. We next derive the stability condition of the steady states by following Ref. [18]. Using the relation 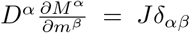 and 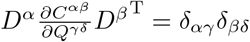, each element of 𝒜 𝒱 is written as

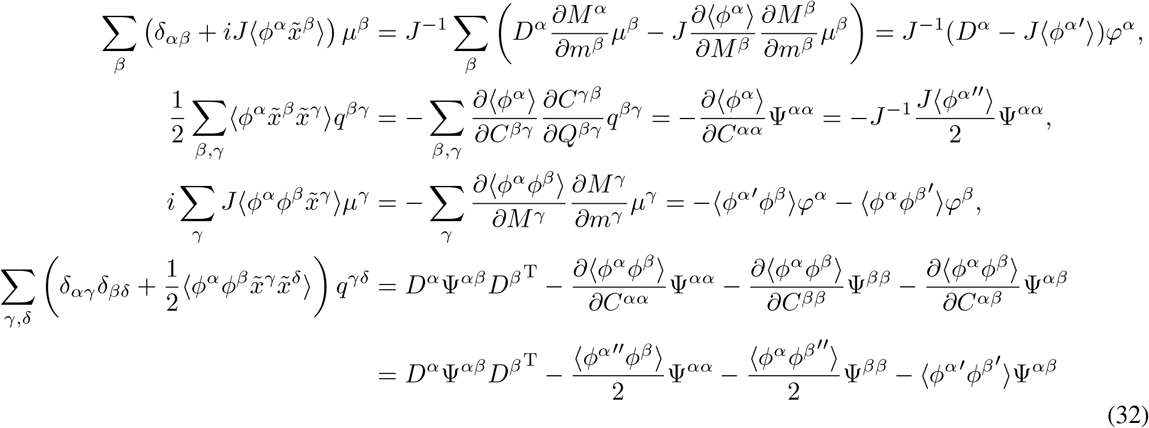

where 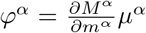and 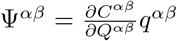, where the operator 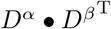acts as 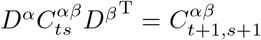 and 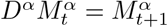.

The steady state is stable if and only if the eigenvalue equation

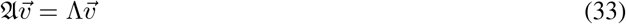

has no solution with the eigenvalue Λ = 0, where the five-dimensional vector 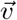 stands for

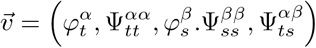

and the operator A in Eq. (33) is given by

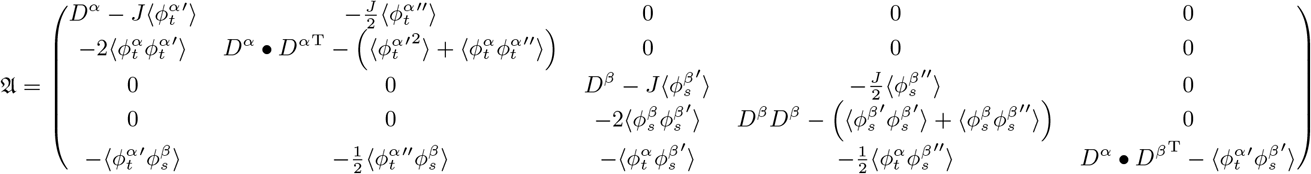

acting onto the vector 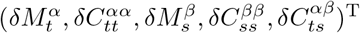.

We first examine the stability of a steady solution, 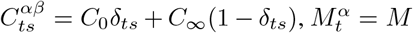. We have to check if there exists a solution to the following equation when Λ = 0

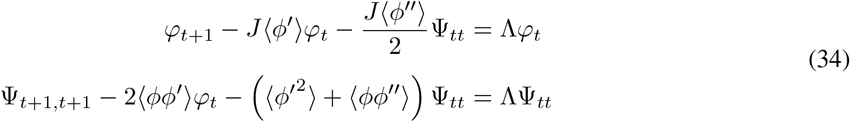

Let the Z−transformation of φ_*t*_ and Ψ_*tt*_ be respectively

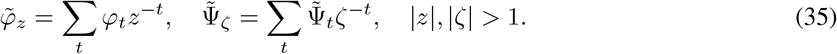

In matrix form, the system of equations (34) can be written as

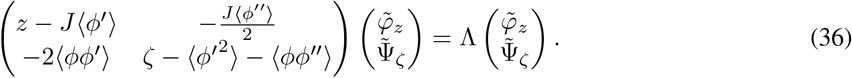

If the eigenvalue Λ = 0 exists, the equation

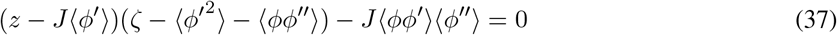

holds true for some z, *ζ* satisfying |z|, |*ζ*| > 1. Now *ϕ* satisfies ⟨*ϕ*^′^⟩ > 0 and ⟨*ϕ*^′2^⟩ + ⟨*ϕϕ*^′′^⟩ > 0; consequently, the steady state is stable against the intra-replica perturbation if

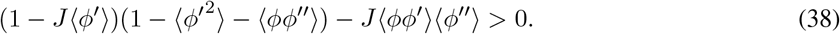

When the steady state is a fixed point (time independent, *C*_∞_ = *C*_0_), the stability criterion within a single replica can be checked by use of Eq. (38). If the steady state is time-dependent *C*_∞_ < *C*_0_, we have to consider the stability against the perturbation 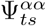. If the inequality (38) holds true, the possibile existence of a vanishing eigenvalue Λ = 0 can be brought about by

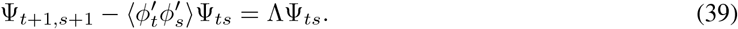

As we saw, the matrix A appearing in Eq. 33 has the form

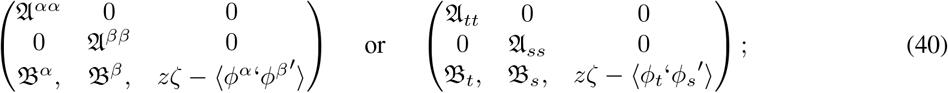

hence, once the stability against the perturbation 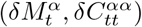 is shown, the instability (possibility of the vanishing eigenvalue) can come from only the terms 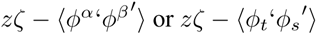respectively.

We next consider the stability against the perturbation 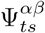, that is, the vector 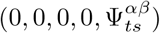. In this case, we have to look at the equation

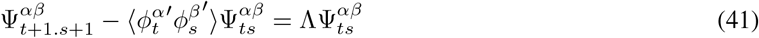

Taking the Z−transformation 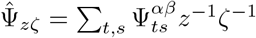 which is defined when |z|, |*ζ*| > 1, we have

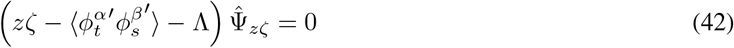

We conclude that The steady state is stable against the inter-replica perturbation 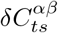, if and only if 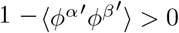 Hence we derive the stability condition

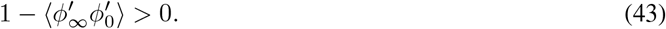

### S4 Derivation of the Formula for the Critical Memory

The meaning of information processing in dynamical systems has become the subject of a vast literature, well summarized in references [36] and [37].

Within reference [36] two possible definitions are given of the memory capacity of a dynamical system. The first one (Eq. 6) does not include any preliminary shifting of mean levels, while the second one (Eq 2.1 of Supplementary Material in Ref [36]) is equivalent to the definition of Ref. [37] and is more natural from the view point of signal processing. An observer in possession of an unbiased estimator for the mean may remove the mean values from all the time series he records; what matters is the relationships between those mean-removed observations and the mean-removed version of the unobserved underlying process. Moreover, we would like the resulting memory capacity to be zero when the linear readout is dominated by a constant baseline value, because nothing can be learned from a readout independent on the input. Adopting therefore the mean-removed formula, we find for the memory capacity *M* in the neighborhood of the second-order phase transition boundary

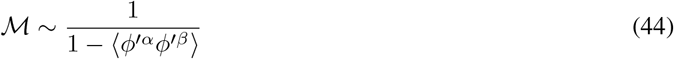

as given in the main text.

To derive this formula, we proceed along the same lines as in Ref. [24], considering the input signal *u*_*t*_ as 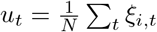 and trying to re-construct the input u(t_0_) with the sparse linear readout 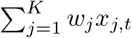 with 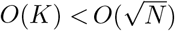. The memory curve *C*_*τ*_ and capacity *C*_*M*_ are given respectively by the determinant coefficient which measures how well the readout neurons reconstruct the past input u(t − τ) correctly, and their sum [37]

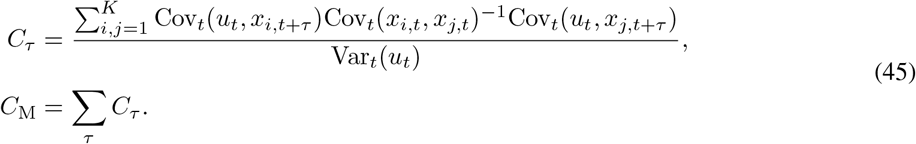

The read out is sparse, so that the covariance Cov_*t*_(*x*_*i*_(*t*), *x*_*j*_(*t*)) becomes diagonal in the infinite population limit *N* → ∞ [23]. Moreover, we deal with the steady state so that this term is constant with respect to time.

We then have to compute 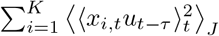. As shown in the Appendix in Ref. [24], when the input signal is a weighted sum of Gaussian random variables, the term ⟨*x*_*i,t*_ *u*_*t*−*τ*_ ⟩ is given by the linear combination of 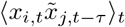, which is the zero-field susceptibility of the parameter 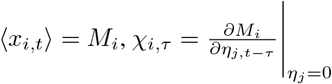

Let the signal be 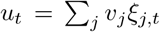. Since we are interested in computing 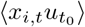, let’s proceed throughout the standard field-theoretical step of inserting an exponential source term for this quantity in side the general functional, to then differentiate by the relevant parameter. The suitable source term is

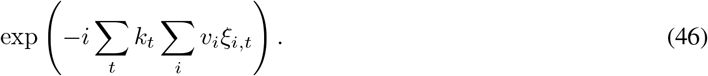

Inserting it into the generating functional, we have

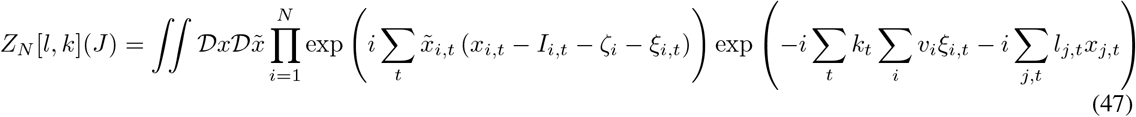

where 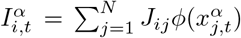, and *ζ*_*i*_ is quenched randomness whose mean and covariance are *µ* and *σ*δ_*ij*_ respectively. Taking average over the dynamical noisy input ξ_*i,t*_ satisfying ⟨ξ_*i,t*_⟩_*ξ*_ = 0 and ⟨ξ_*i,t*_ξ_*j,s*_⟩_*ξ*_ = *σ*_in_δ_*ij*_δ_*ts*_ we have

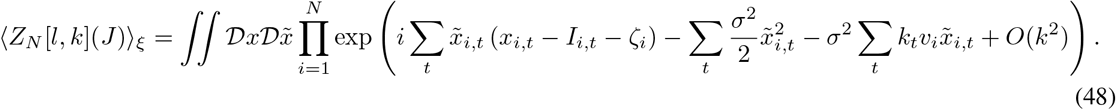

Thus, the term 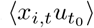 is found to be given by the weighted sum of the linear responses 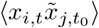 as

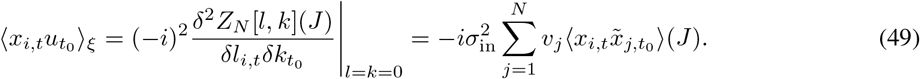

The next quantity needed is 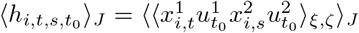. To compute this, we insert in the generating functional the single source

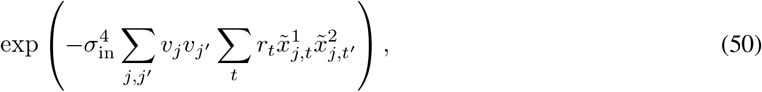

This turns the the generating functional into

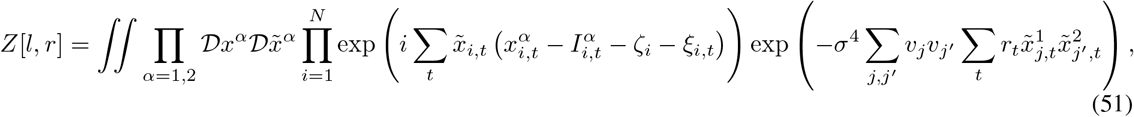

which makes additional perturbation to the inter-replica correlation (see Eq. (G11) in Ref. [24]).

Let v_*i*_ be ∼ 1/N. The form of r_*t*_ is assumed to be 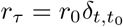. Using it, 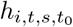 is written as

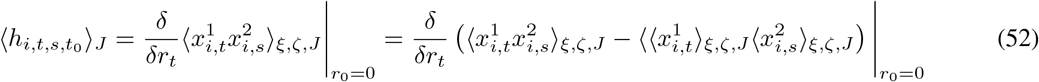

The last equality is brought about by 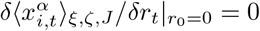 (which is derived through Wick’s theorem [42] due to *x*_*i,t*_ being Gaussian random variables when we take the infinite population limit *N* → ∞) and by use of the causality or normalization condition which gives 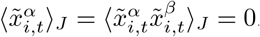.

Further, it should be noted that 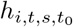 is a perturbation brought about by the additional source term (50), so that

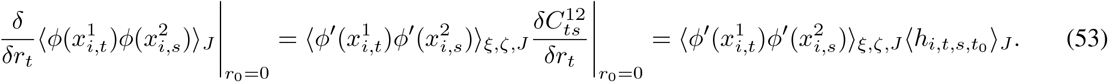

Let 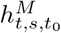 be

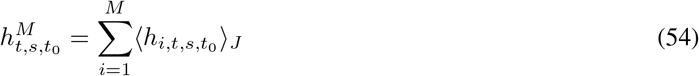

for *M* = 1, 2 …, N. What we desire is 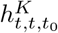, which satisfies

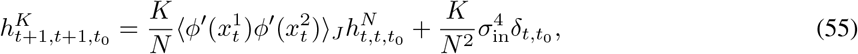

where the last term is coming from the random inputs

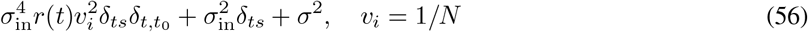

The term 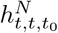 in the left hand side evolves as

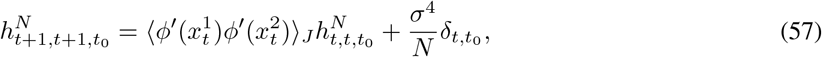

so that we have, in the steady state,

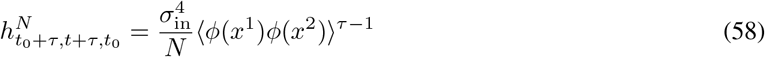

and further, we have

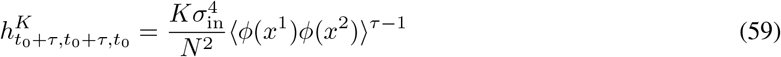

from Eq. (55).

The memory curve *C*_*τ*_ is proportional to 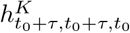, so that we conclude that the capacity 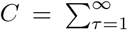 satisfies

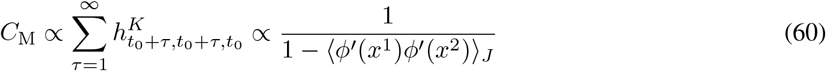

